# Dissolved inorganic carbon driven dynamics of calcite shell formation in 12 strains of the freshwater algae *Phacotus lenticularis* (Chlorophyta)

**DOI:** 10.1101/2025.05.04.652085

**Authors:** Uta Gruenert, Jan Benda, Oliver Bossdorf, Uta Raeder

## Abstract

This article explores the close relationship between dissolved inorganic carbonate ion concentration (DIC) and the calcification state of *Phacotus lenticularis*, a globally abundant freshwater phytoplankton that is responsible for a significant part of modern lake carbonate production during bloom formation. We cultured 12 freshly isolated *P. lenticularis* strains under an ecologically relevant range of DIC (0.2 to 12 mmol l^-1^ total scale) for 14 days. From this experiment we gained information on responses in shell formation and growth that highlight strong lower boundaries in morphometric calcite shell variables with regards to DIC. All *P. lenticularis* strains showed reduced shell thickness by up to 60 % and dissolved calcite crystals structures at declining DIC < 4 mmol l^-1^. Increasing DIC > 4 mmol l^-1^ had no significant effect on shell thickness and crystal length in the culture experiments. We found a significant preadaptation of all 12 strains to ambient DIC concentrations measured in their lake of origin, but no dependence of growth rates up to a lethal DIC of > 10 mmol l^-1^. The simulation experiments illustrate the close relationship between shell function and dissolved inorganic carbonate ion concentration in lakes and highlight the need of continued research of important roles in biogenic carbon transformation and storage in a future world.

## Introduction

Biological calcification is an important climatic feedback, and in phytoplankton is highly impacted by changes in water carbonate chemistry (Langer et al. 2009, Müller et al., 2010, Rokitta and Rost 2012, Gruenert and Raeder 2014, Raven and Beardall 2021, Page et al. 2022). In alkaline lakes, which comprise more than half of all inland water, dissolved inorganic carbon (DIC) is the dominant form of aquatic carbon (Khan et al. 2020). DIC in water is composed of bicarbonate (HCO_3−_) and carbonate ions (CO_32 −_). These anions are associated with gaseous carbon (free CO2) via the carbonate equilibria. DIC levels in lake water vary considerably on seasonal and daily scales with values between 0.1 to 100 mmol l^-1^ (Cole 2013, Song et al. 2018). Variability depends primarily on carbonate inputs (riverine and groundwater loads, atmospheric CO_2_ input) and carbonate outputs (water outflow, ground-water leakage, degassing of CO_2_ to the atmosphere, calcite precipitation (Hammer 1986, Finlay et al. 2009, Müller et al. 2016, He et al. 2022). Importance of lake metabolism in driving DIC changes has been found to diminish with increasing alkalinity (Marcé et al. 2015, Khan et al. 2020). Various aspects of climate change, including alterations in precipitation regime promoting higher weathering rates and ion transfer, falling lake and groundwater tables, as well as increased soil respiration have been identified as relevant drivers of hard-water lake chemistry (Battin et al. 2009, Tranvik et al. 2009, Li et al, 2013, Rogora 2020).

Future trends for these lakes remain uncertain and predict increasing ion concentrations and greater chemical variation (Rinta et al. 2015, Weyhenmeyer et al. 2024). While growth of freshwater phytoplankton is found to be stimulates in future climate change scenarios it remains speculative how calcite production and competitive fitness of calcifying phytoplankton will be effected (Shi et al. 2017).

The most abundant modern species of freshwater calcifying phytoplankton is *Phacotus lenticularis*, a bloom forming member of the order *Chlamydomonadales* which typically have two flagella for locomotion (Grünert et al. 2016, Lenz et al. 2018). The unicellular alga fixes dissolved inorganic carbon by forming a lens-shaped calcite shell. Shell formation is thought to be genetically controlled. Ions are concentrated inside intra-cellular vesicles to form small calcite crystals. The crystals exit the cell and fuse to form a thin continuous shell that covers the outer lorica surface (Shaked et al. 2023). A second phase of growth subsequently forms the highly organized ring-like calcite crystals structure of the mature shell. It is yet unknown if *P. lenticularis* uses specific ion channels and pumps in the daughter cell plasma membranes to increase Ca^2+^ concentrations and pH. Only two other unicellular marine calcifiers, coccolithophores and foraminifera, have been found to perform intracellular crystal formation (de Nooijer et al. 2009, Mackinder et al. 2010).

*P. lenticularis* shells consist of 98% – 99% pure CaCO_3_ and contribute significantly to the removal of epilimnetic carbon via the combined effects of calcification and the downward transport of organic carbon in sinking aggregates to the sediments (Krienitz et al. 1993, Klaas and Archer 2002, Gruenert and Raeder 2014, Lenz et al. 2018). At the same time, the precipitation of calcium carbonate during calcite shell formation releases CO_2_ and this way reduces total alkalinity (Gilbert et al. 2022). *P. lenticularis* influences carbonate flux in hardwater lakes, particularly during bloom periods (Gruenert et al. 2016, Lenz et al. 2018). Lake sediments containing significant amounts of *P. lenticularis* shells were reported from many continents (Burns and Mitchel 1974, Haberzettl et al. 2007, Lenz et al. 2020).

A growing number of studies on marine pelagic coccolithophore species, strains and morphotypes suggests variable response patterns of calcification rates, shell morphology and growth to changes in carbonate chemistry in particular to elevated CO_2_ partial pressures (Langer et al. 2009, Müller et al. 2015, Feng et al. 2017, Kottmeier et al. 2022). These findings have been linked to a high genetic variability in coccolithophores due to their global distribution (Medlin et al. 1996, Iglesias-Rodriguez et al. 2006). Photosynthesis and calcification of *Emiliania huxleyi* were stimulated due to additional HCO_3_^-^ uptake in a high DIC concentrations culture experiment, whereas growth was unaffected (Kottmeier et al. 2016). The same experiment showed inhibited coccolithophorid growth and calcification at increased H^+^ concentrations. Adaptation by increased tolerance to ocean accidification has been observed in experimental evolutionary studies on clonal replicates of coccolithophores (Chan et al. 2021). Zhang et al. (2018) additionally found higher optimal pCO_2_ growth and higher tolerance to low pH in populations isolated from places with larger environmental variability. The work to date is already providing valuable information on marine coccolithophores sensitivity or resilience to climate change. Still so far, there has been no systematic examination of how projected changes in carbonate chemistry will affect the physiology and calcification of freshwater calcifying phytoplankton.

How DIC influences calcification of freshwater phytoplankton is a central component for understanding mechanisms of biological carbonate formation in lakes and their relationship with climate change. We fill this gap by performing trait based culture experiments on freshly isolated *P. lenticularis* wildtype strains over a wide DIC range. The spatial isolation of alkaline lakes can limit dispersal of freshwater phytoplankton, restrict gene flow between geographic populations, and result in their genetic differentiation (Van Den Wyngeart et al. 2015, Rengefors et al. 2017, Ryderheim and Kiørboe 2024). Freshwater calcifying algae have rarely been studied because of difficulties to isolate and cultivate this algal group for controlled experiments. Furthermore, it is often argued that experiments using long-term laboratory cultures may document evolutionary adaptations to changes induced by the culture conditions (Zhang et al. 2021). We isolated 12 strains from two lakes in South Germany differing in lake morphology and DIC content and immediately assessed their growth, cumulative inorganic carbon production and calcite shell morphology parameters as a function of DIC and carbon ion concentrations. Our study aims to investigate whether geographical variation in a lakes ambient carbonate chemistry exerts a selection pressure on *P. lenticularis* shell formation.

## Methods

### Sample collection

Plankton samples were collected by 100 μm mesh net tows taken from lake Gönningersee (48°25’32.5”N 9°10’38.4”E) and lake Großer Ostersee (47°47’30.6”N 11°18’03.5”E), South Germany between the month July and September. Lake Gönningersee is a shallow (maximum depth: 4 m), mesotrophic lake receiving high but variable input of carbonate rich stream water. Lake Großer Ostersee is a deep (maximum depth: 29 m), oligotrophic lake with relatively little variation in pH and DIC and regular groundwater inflow of carbonates. Seasonal DIC concentrations in the lake surface water ranged from 4.8 to 5.7 mmol l^-1^ in lake Gönningersee and from 4.2 to 4.7 mmol l^-1^ in lake Großer Ostersee. Environmental parameters for the sampling locations were measured directly before plankton sampling (temperature and pH) and in vitro (total alkalinity (T_A_) and major nutrients).

*Phacotus lenticularis* cells were isolated from plankton samples by serial dilution with specific microcapillaries (borosilicate; 1.5-mm outer diameter; GB150F-8P; Science Products, Hofheim, Germany) pulled on a P97 puller (Sutter Instruments, Novado, CA, USA) and broken to an appropriate diameter. N-HS-Ca culture medium was prepared as described in Schlegel (2000). Nitrate (NO_3_ ^-^), phosphate (PO_4_ ^3-^) and calcium (Ca^2+^) were added in concentrations representing ambient lake nutrient concentrations of 160, 10 and 1500 µmol l^-1^, respectively. *P. lenticularis* cells were placed in culture medium immediately after isolation. Strains from lake Gönningersee likely belong to morphotype III while strains from lake Ostersee belong to morphotype IV as distinguished by Schlegel (2000).

### Experimental Design

*Phacotus lenticularis* strains were inoculated in triplicate shake flasks and assigned to a range of DIC concentrations (7 treatments, DIC = 0.2, 1, 4, 6, 8, 10, 12 mmol l^-1^). The lower DIC levels corresponded to seasonal variations found in middle and hard water lakes (0.2 – 6 mmol l^-1^, Stumm & Morgan, 2012). The higher DIC levels exceeded this range up to the point where no growth of *Phacotus* cells was monitored anymore. Treatment culture medium was prepared using particulate and dissolved inorganic carbon free N-HS-Ca culture medium. DIC and T_A_ were increased by addition of sodium bicarbonate and magnesium carbonate. Carbonate chemistry manipulation is more analogous to changes expected in lake surface waters in the next decades than gaseous CO_2_ addition. Cultures were grown at 20 °C (±0.15 °C) with an incident photon flux of 100 µmol photons m^-2^ s^-1^ PAR (ATUM LEDbar 18W, Klutronic GmbH, Klagenfurt) under a stable 12 hour light/dark cycle. T_A_, DIC, pH and temperature were measured immediately after preparation of the medium, at the beginning of the experiment, after 7 days and after 14 days at the end of the experimental treatment to control for stable DIC and T_A_ concentrations. Calcium concentrations were kept constant at 1.5 mmol l^-1^ and pH was not manipulated.

Alkalinity was determined by acidimetric titration with 0.01 M HCl, in triplicates (Dickson et al., 2007). DIC concentrations were calculated from measurements of acid neutralising capacity, pH, temperature, and specific conductivity according to Mackereth et. al (1978) and reported as the average and standard deviations of triplicate measurements. The relative uncertainty for both DIC and T_A_ was ±0.5 % of the final value. Temperature and pH were measured using a glass electrode (Xylem, pH535, Weilheim, Germany), which included a temperature sensor and was two-point calibrated with NBS buffers prior to every set of measurements. Average repeatability was found to be ±0.03 pH units. Conductivity was measured using a Conductivity/TDS/Salinity Meter (Extech EC400). Ion exchange chromatography was used to obtain concentrations of inorganic ions in solution (Ca^2+^, Mg^2+^, K^+^, Na^+^, Cl^-^, F^-^, NO_3_ ^-^, PO_4_ ^3-^ and SO_4_ ^2-^) (Dionex Dx-120, Thermo Fisher Scientific, USA, California). The carbonate system was calculated from direct measurement of pH, temperature, T_A_ and concentrations of primary ions using the software WinIAP (Sequentix, http://www.sequentix.de/software_winiap.php). Activity coefficients were estimated by means of the extended Debye–Hückel equation, valid for solutions of higher ionic strength and electrical conductivities (Stumm and Morgan, 1996).

Strains were acclimated for 24 hours in dilute batch culture for each experimental condition prior to inoculation. Cells were inoculated in treatment medium at an initial cell concentration of ∼ 50 cell ml^-1^ in 500 ml shake flasks and placed on a shaker. In the 4, 6 and 8 mmol l^-1^ DIC treatments, where higher cell densities of approx. 40 000 cells ml^-1^ were reached, 10 ml aliquot was sub-cultured into new medium after seven days to minimize changes in carbonate chemistry and nutrient availability and to avoid self-shading. Cultures were kept at experimental DIC concentrations for 14 days.

### Analysis

Plastic responses of functional morphological traits were measured as (i) shell diameter and thickness, (ii) shell ultra-structure, (iii) bulk calcification rates for each strain of *Phacotus* cells, as well as (iv) cell concentrations and (v) growth rates. Subsamples for measurements of cell concentrations were taken every second day and placed on a glass microscope slide (improved Neubauer haemocytometer). Four images per strain and treatment were taken using a light microscope with digital camera. By means of the ImageJ software package the images were converted to 8-bit images and cells were segmented from the image background. Cell counts as well as cell sizes were retrieved using the particle analyzer algorithm. Cell size was reported as the average and standard deviation of measurements retrieved from four images.

Cell densities were calculated by dividing the cell counts by the measurement volume of 0.1 µl. For cell cultures that have been diluted after 7 days, all cell densities after dilution were multiplied by (1−Δ*V/V)*^*-1*^=4, where V=40 ml is the volume of the treatment medium and Δ*V=*30 ml is the removed volume, in order to correct for the dilution. Specific growth rates μ were obtained via a logistic function

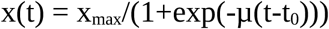

that has been fitted to the cell densities *x* measured at times *t* (=1, 3, 5 … days). *x*_*max*_ is the maximum cell density reached at saturation and *t*_*0*_ is a shift parameter.

Samples for scanning electron microscope (SEM) were taken during exponential growth from the optimum as well as the lower and upper boundaries of the growth response functions (DIC 1, 4 and 8 mmol l^-1^) and placed onto wafers, air-dried at room temperature for 24 hours, then sputter coated with gold-palladium. Imaging was carried out with a EVO LS10 SEM (ZEISS, Oberkochen, Germany) at an acceleration voltage of 20 kV. Calcite shell morphological traits were measured as (i) shell diameter, (ii) shell thickness, (iii) pore size, and (iv) calcite crystal length for each *Phacotus* strain (total number of images N=300). Shell thickness was measured as rim width from images showing shell halves. Calcite crystal length was estimated from shell surface images as follows. First, images showing the lens-shaped shell from the top were cropped to a square focusing on the shell’s center (Fig.1A), transformed to gray-scale, and slightly smoothed using a Gaussian filter with a sigma of one pixel. The lens-shaped shell surface caused a gradual change of the mean brightness towards the rim. We corrected for this by fitting and then subtracting a 2-dimensional parabola from the image. The crystals are characterized by particularly bright or dark edges resulting in prominent peaks and troughs in brightness along a section through the image. In each row and column of pixels we detected these peaks and troughs (Fig.1B), computed a typical crystal length by dividing the image width by the averaged number of peaks and troughs, and calculated the median. All shell measurements are reported as means with standard deviations per DIC treatment and lake.

**Fig. 1.**
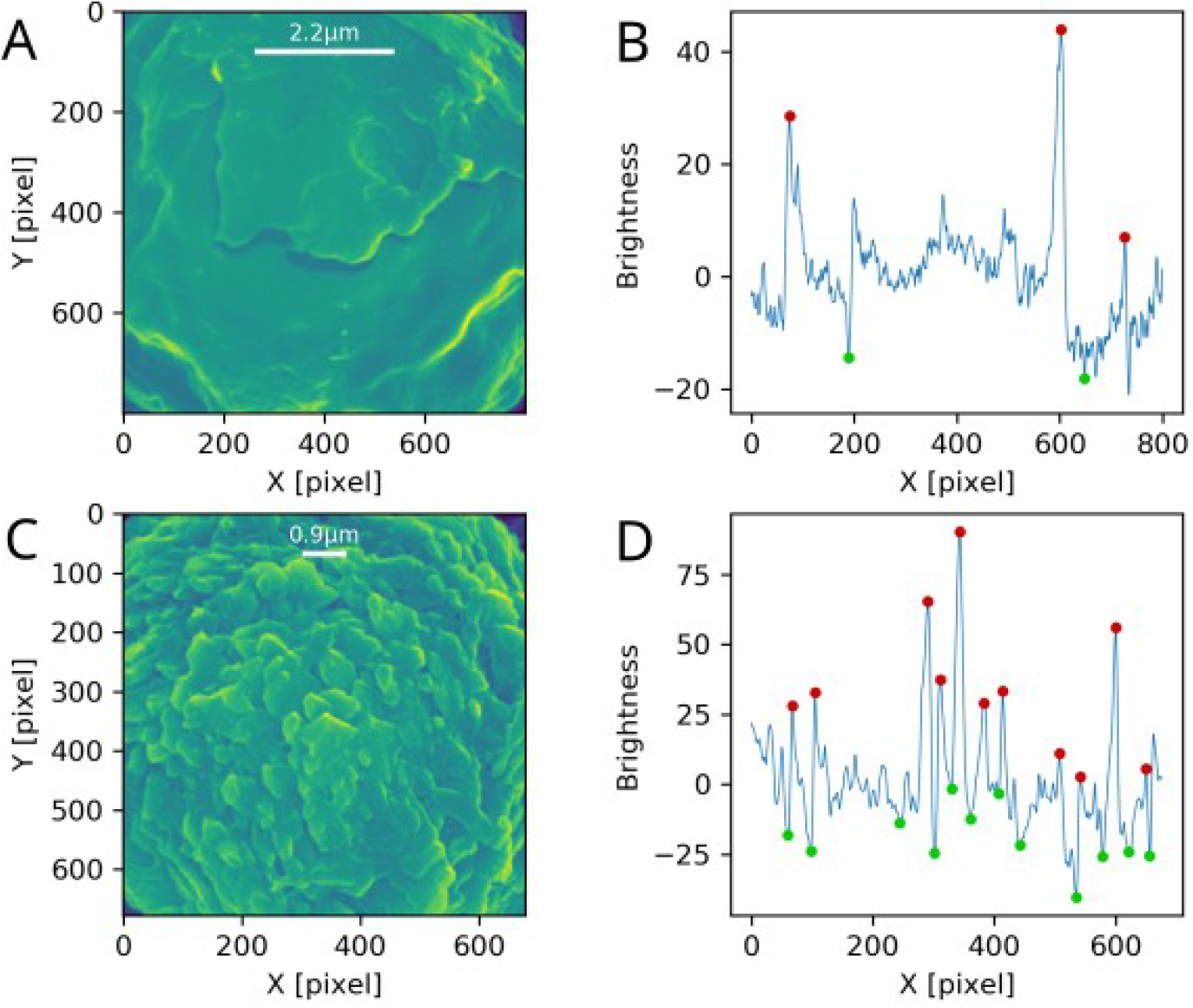
Smoothed gray-scale shell surface images showing bright or dark edges of individual crystals A,C; Peaks and troughs of brightness were detected along a section through the image B,D

### Statistical analysis

We used least square regression and Pearsons correlation coefficient for linear correlations and nonlinear least squares functions for nonlinear models to explore relationships between environmental parameters (T_A_, pH, HCO_3_^-^, CO_3_^2-^).

An exponential function with a rate constant set to the average of the calculated growth rates λi was then fitted to the cell counts in order to calculate exponential growth (Gruenert et al. 2014). From final cell densities at the end of the treatment position, height, and width of the optimum peak were estimated by least-square fitting of a parameterized Gaussian function to the data across all DIC concentrations. The functions were selected based on fit quality measures (correlation between fit and data, root-mean-square difference, Gaussian distribution of residuals). Differences of response curves between the different *P. lenticularis* lineages and DIC concentrations were assessed based on confidence intervals of the fitted curves obtained by bootstrapping. Differences in the parameters obtained from the fits, like peak position and width of the optima, were tested by a one-way ANOVA with the strains, lake origin, and DIC concentration as factors. Test were performed on shell characteristics (shell diameter, shell thickness, calcite crystal size and pore size) to evaluate differences in all strains. Furthermore, principal component analysis (PCA) of the fit parameters and data were used to quantitatively explore patterns within the algal strains response functions. All calculations and the ordination biplots were made using R Version 3.4.3.

## Results

The DIC culturing system (DIC gradient between 0.2 to 12 mmol l^-1^) proved effective at maintaining a steady carbonate system in treatment replicates over the course of the experiment. DIC and total alkalinity (T_A_) depletion by the physiological activity of *Phacotus* cells was higher in low DIC treatments but never exceeded 6 % of the initial concentrations in the culture medium. The calcite saturation state (Ω_Calcite_) was 0.5 at DIC 1 mmol l^-1^, 9.3 at DIC 4 and 51 mmol l^-1^ at DIC 8 mmol l^-1^. As expected, alkalinity was proportional to DIC with a proportionality factor of 0.97 meq/mol (Fig.2A). In the culture medium we did not control for pH. Values of pH increased quickly from 6.2 to approx. 7.3 for DIC values < 1 mmol l^-1^. For larger DIC values pH further increased approx. linearly up to 9.2 (Fig.2B). HCO_3_ ^-^ was almost linearly related to DIC (Fig.2C). CO_3_ ^2+^ concentrations were about an order of magnitude smaller than HCO_3_ ^-^ and increased according to a power law with exponent 2.5 with DIC (Fig.2D). The concentration changes of both CO_3_ ^2+^ and HCO_3_ ^-^ follow the expected dependency on pH.

**Fig. 2.**
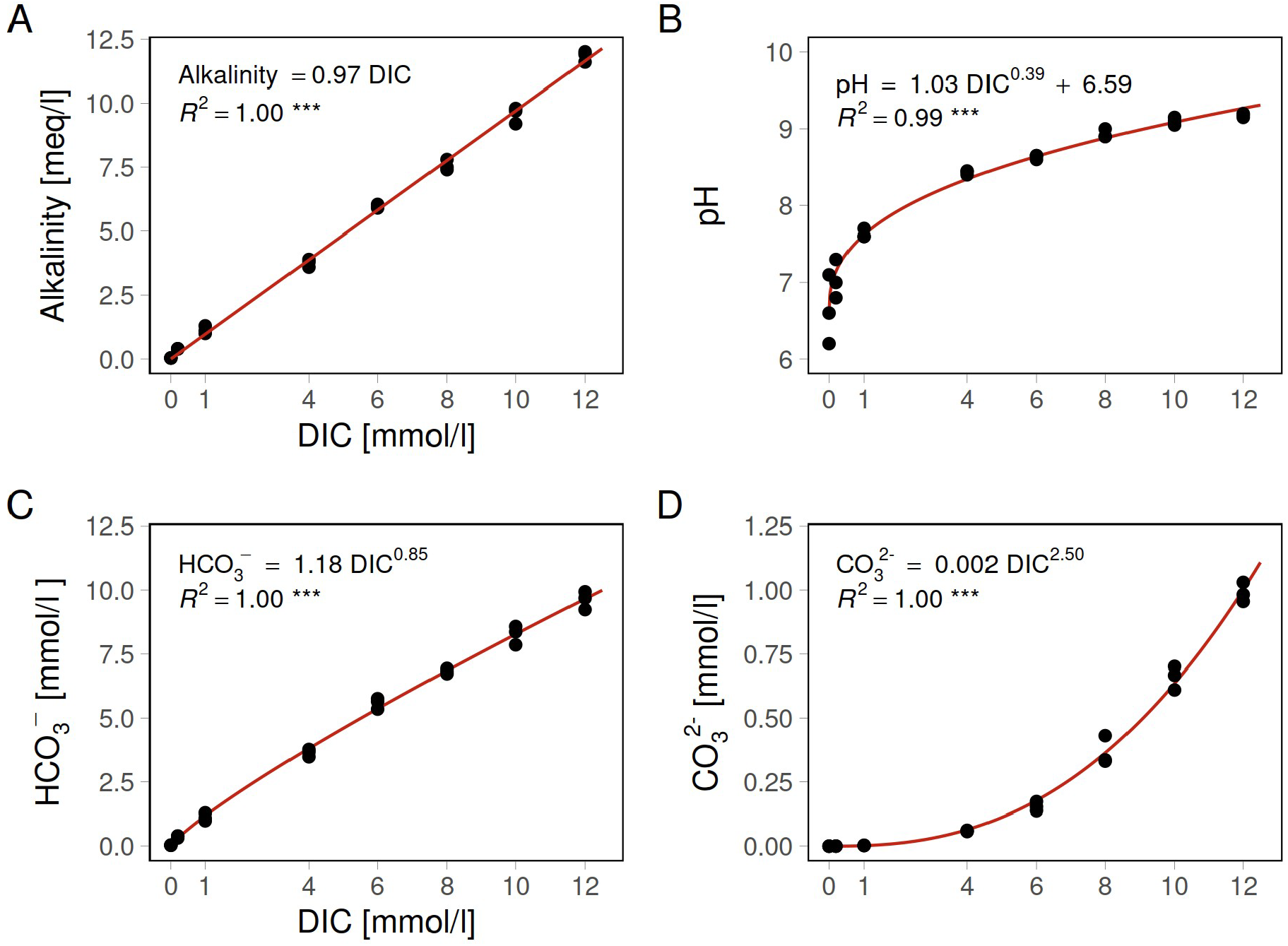
Non-linear and linear relationships between Alkalinity (A), pH (B), HCO_3_^-^ (C), CO_3_^2-^ (D) and DIC concentrations in the modified culture medium N-HS-Ca. Coefficient of determination R^2^ as a measure of the goodness of fit and significance levels are presented for each regression. Curves were fitted based on nonlinear least squares functions for nonlinear models and least square regression and Pearsons correlation coefficient for linear correlations.

### Effect of DIC concentrations on shell characteristics

We measured shell morphology parameters of 300 complete *P. lenticularis* calcite shells and 170 shell halves based on SEM observations at three DIC concentrations (1, 4 and 8 mmol l^-1^). The images provided high resolution information about shell diameter, thickness, shape and arrangement of crystals, as well as pore size (Fig.4). Measured average diameter of *Phacotus* strains was 12.0±1.9 µm in OS strains across DIC concentrations of 1, 4 and 8 mmol l^-1^ and was significantly smaller than average diameter of GS strains (13.6±1.8 µm) (Wilcoxon rank test, W=8092, p<0.0001). This corresponds well with SEM image measurements of *Phacotus* shell diameters from both lakes (Fig. 3 and Lenz et al. 2018). Shell diameters were largest at ambient DIC values of 4 mmol l^-1^ and decreased by 1.5 and 1.2 µm when DIC was changed to 1 and 8 mmol l^-1^, respectively (Fig. 4A). A large decrease in shell thickness was found when DIC concentrations were lowered from 4 mmol l^-1^ to 1 mmol l^-1^ in all strains (Wilcoxon rank test, W=590, p<0.0001). In strains isolated from lake OS shell thickness decreased from 1.7±0.3 µm to 0.6±0.2 µm, while GS strains showed a smaller reduction from 1.8±0.3 µm to 1.2±0.2 µm. In contrast, shell-thickness of *P. lenticularis* strains did not change noticeably when DIC concentration was doubled to 8 mmol l^-1^ (Fig. 4B).

**Fig. 3.**
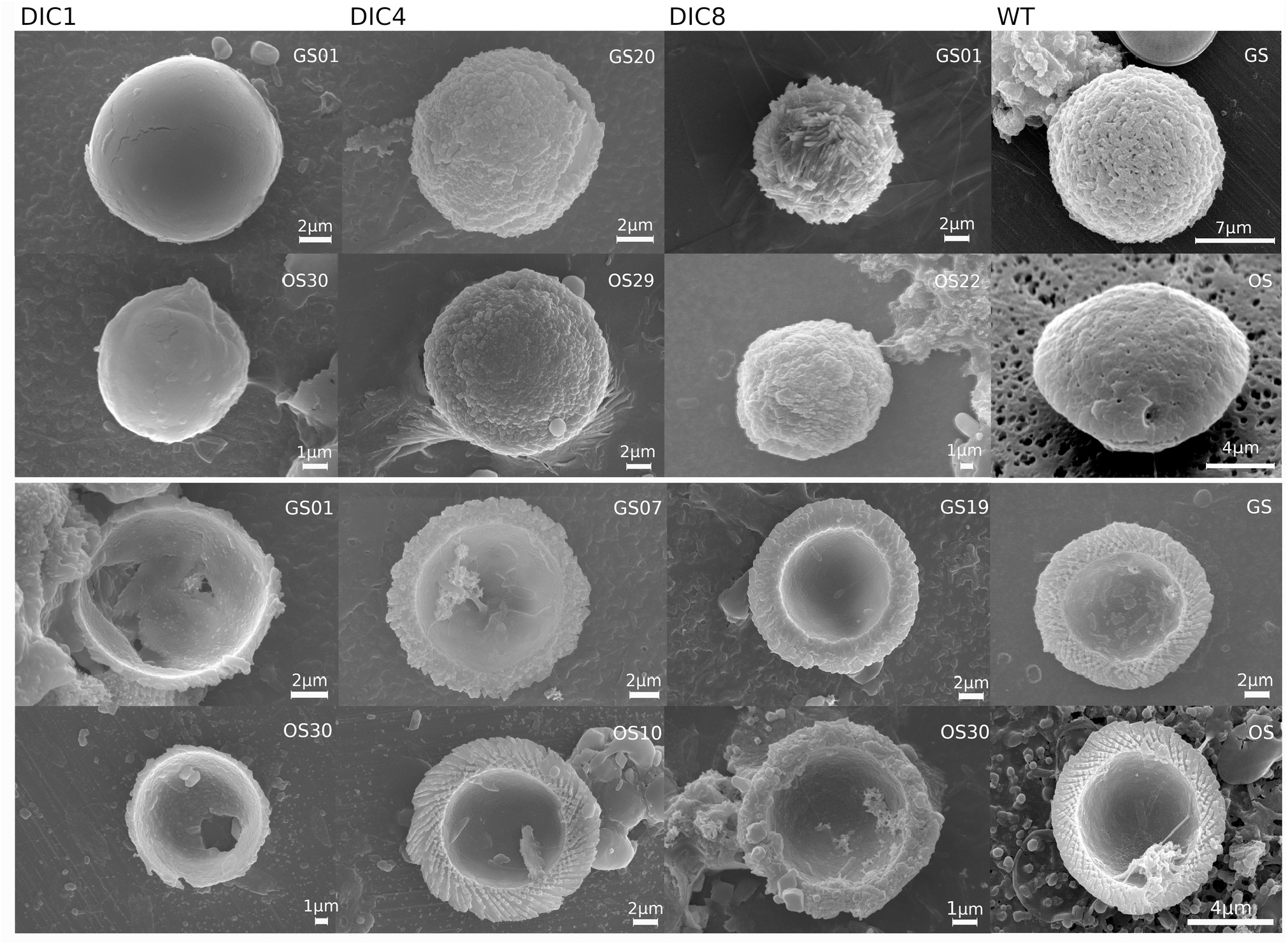
Scanning electron micrographs showing open shell halves (upper panel) and complete shells (lower panel) of *P. lenticularis* strains across three DIC concentrations; calcite shells are composed of two shell halves attached to one another at the rhim. The most rightern column WT displays original wildtype *Phacotus* shells from lake Gönningersee (GS) and lake Großer Ostersee (OS). Strain IDs are shown for each image.

**Fig. 4.**
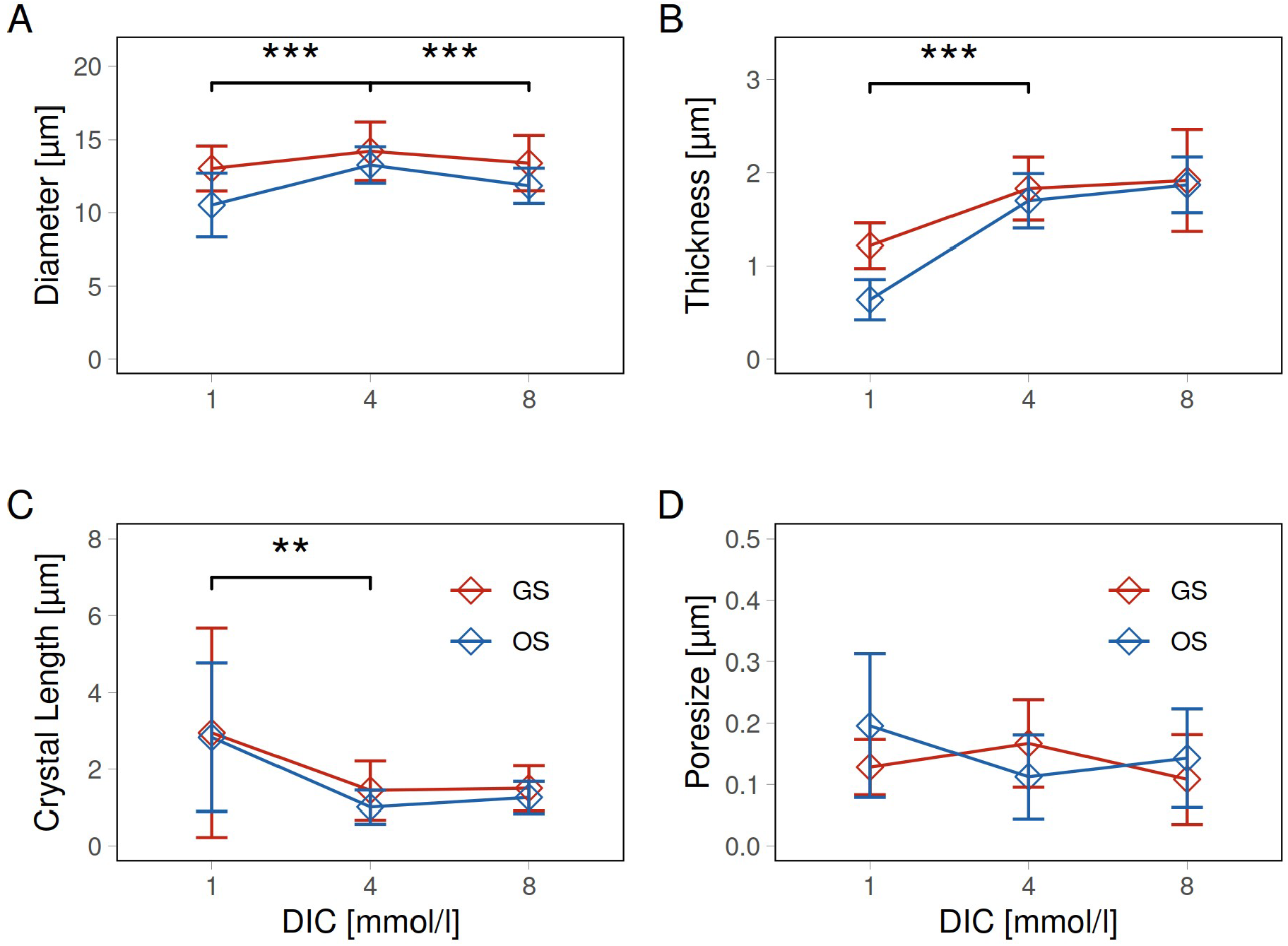
Calcite shell parameters versus DIC for *P. lenticularis* strains per lake; diameter (A), thickness (B), crystal length (C) and poresize (D) were measured with the aid of a scanning electron microscope, vertical error bars represent standard deviations of between 20 and 40 measurements for each strain isolated from lake Gönningersee (GS) and Grosser Ostersee(OS).

Shells exposed to a low DIC concentration of 1 mmol l^-1^ lost their distinct crystalline structure with the calcite surface becoming smoother and more fragile (Fig.3). Adjacent crystals merged to a larger calcite field accompanied by a loss of crystal thickness. This is reflected by a significant increase in average crystal length from 1.0±0.4 µm to 2.8±1.9 µm in OS strains and from 1.4±0.7 µm to 3.0±2.0 µm in GS strains (Fig. 4C, Wilcoxon rank test, W=2235, p<0.0001). An increase in DIC from 4 to 8 mmol l^-1^, however, had no significant effect on average crystal length. Ambient crystal length was preserved despite a high tendency for CaCO_3_ precipitation when DIC was doubled. Our SEM image analysis, however, demonstrated lake-specific differences in crystal shape when DIC increased from 4 to 8 mmol l^-1^. GS strains largely belong to morphotype III defined by a rhombohedral crystalline shape with a high density of nanoscale pores at ambient DIC. OS strains belong to morphotype IV defined by a smooth shell surface and lower pore density. While almost all shells lost their crystalline structure at DIC 1, the crystals of GS strain shells elongated to spines that extended outward at a DIC of 8 mmol l^-1^ (Fig.5). Calcite crystals of OS strain shells developed a rhombohedral shape at DIC of 8 mmol l^-1^, similar to the shells of GS strains at ambient DIC concentrations of 4 mmol l^-1^ .

**Fig. 5.**
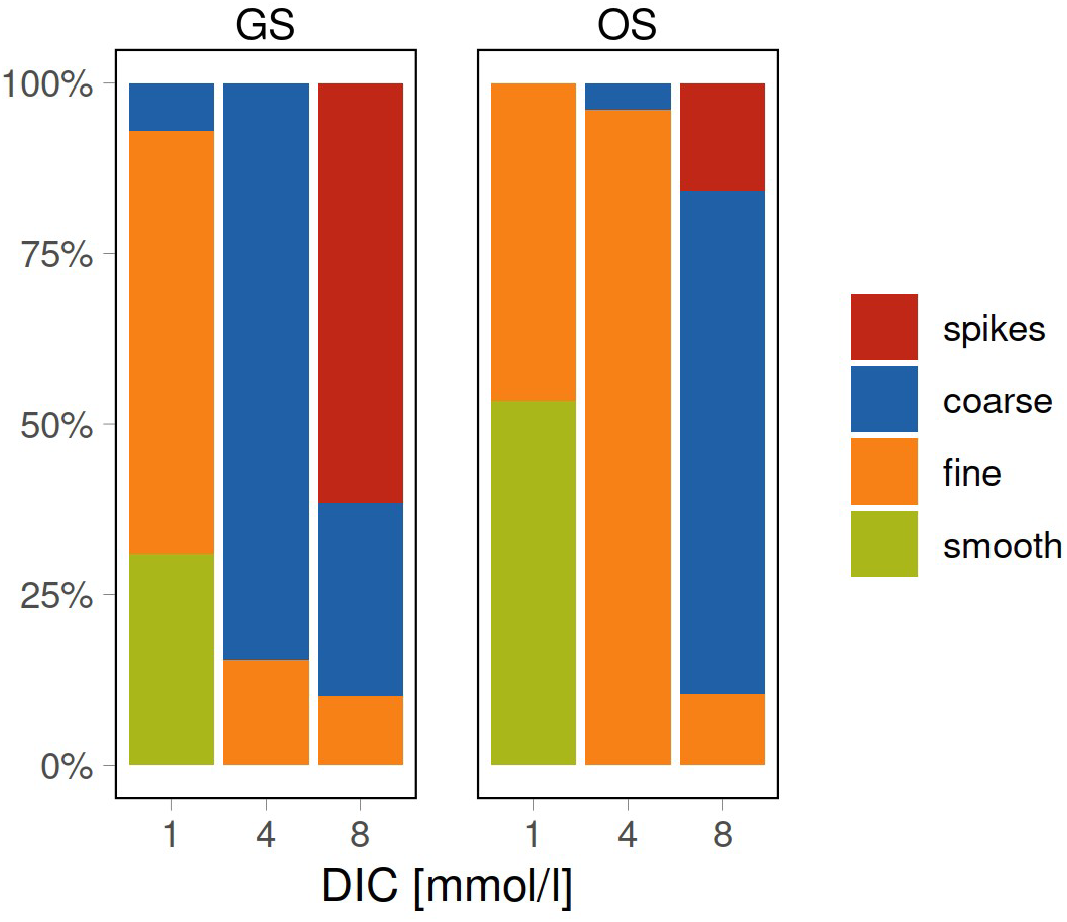
Percent stacked bar plots representing the proportions of four categories of calcite crystal shape at the surface of *P. lenticularis* shells across three DIC concentrations comparing lakes Gönningersee (GS) and Großer Ostersee (OS). Crystal shape estimates were obtained from SEM pictures of *P. lenticularis* shells.

Pore size of *Phacotus* shells did not change significantly between treatments (Fig.4D). Surprisingly, many shells lacked pores entirely at all three DIC concentrations with no visible effect on cell growth (see below). A small space between the rims where the two shell halves attach to one another may allow for a selective uptake of molecules from the surrounding medium when pores are absent.

### Growth rate and cell density

The 12 *P. lenticularis* strains did not alter their growth rate in response to DIC manipulations across a broad range between 0.2 and 10 mmol l^-1^ (Fig. 6A,B). Estimated growth rates did not differ between OS strains (0.49 ± 0.15 d^-1^) and GS strains (0.55 ± 0.14 d^-1^, t-test, t=0.53, p=0.60). At a concentration of 12 mmol l^-1^ all strains except GS20 died within a few days into the experiment.

**Fig. 6.**
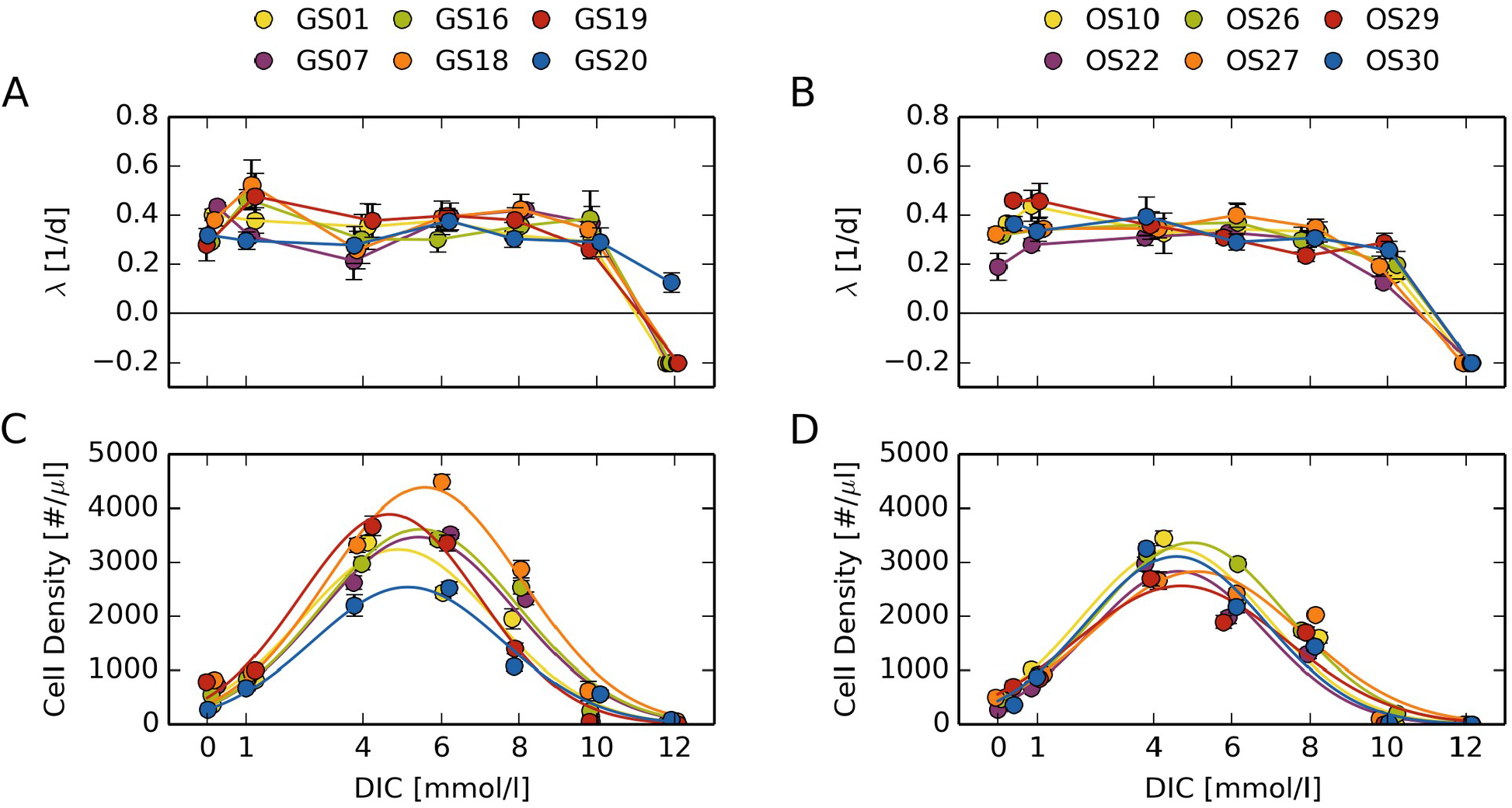
Optimum curve responses of measured growth (λ) and cell density at the final day of the experiment (14 days) across a DIC gradient, individual *P. lenticularis* strains from lake Gönningersee (left) and lake Großer Ostersee (rigth) are plotted.

In contrast to growth rates, maximum cell densities of *P. lenticularis* strains strongly depended on DIC concentrations (Fig.6 C,D). Cell densities measured at day 14 followed a Gaussian response profile. At low and high DIC concentrations cell cultures reached their stationary phase earlier in the experiment, resulting in a strongly reduced cell density compared to the one at ambient DIC concentrations. However, based on maximum cell counts, all 12 strains showed a wide tolerance to DIC concentrations between 1 and 8 mmol l^-1^. Peak height (F-test, F=3.5, p=0.09) and width (F-test, F=0.002, p=0.97) of the Gaussian functions fitted to the cell densities did not differ between lakes. GS strains, however, preferred slightly higher DIC concentrations than OS strains (F-test, F=6.8, p=0.03). Optimum growth of GS strains occurred at DIC concentrations of 5.2 ± 0.3 mmol l^-1^ and of OS strains at 4.7 ± 0.2 mmol l^-1^, matching ambient lake concentrations and provided evidence for local adaptation to their lake of origin (Fig.7). Observed strain specific preferences did reflect environmental DIC and pH conditions in their lakes of origin, as indicated by results from reciprocal crosses. Further, variability in peak height of the Gaussian preference function was larger in GS strains than in OS strains (F-test, F=7.4, p=0.046, Fig.6 C,D). Variable thresholds either could result from strain-specific differences or could be altered by culture conditions.

**Fig. 7.**
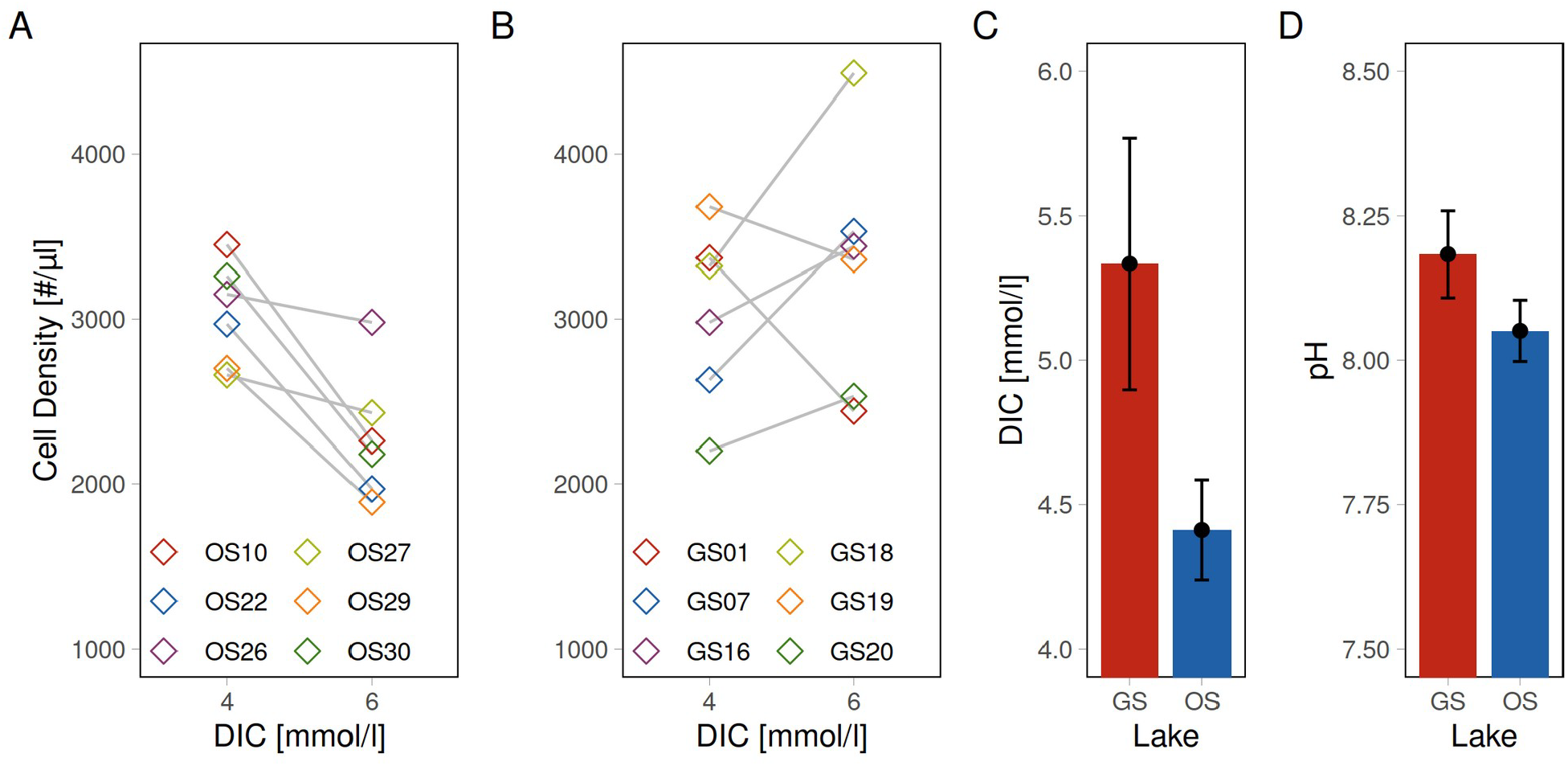
Results from reciprocal crosses show response of cell density at the final day of the experiment for individual *P. lenticularis* strains at two DIC concentration representing ambient conditions in the two lakes of strain origin (A, B); average DIC concentration with standard deviation (C), average pH and standard deviation during the vegetation period (C) in lake Gönningersee (GS) and lake Großer Ostersee (OS).

The *P. lenticularis* strains from the two lakes formed two distinct groups along the first principal component of the multivariate space spanned by all their characteristics (Wilcoxon test, W=0, p=0.002). GS strains differed from OS strains by larger shell diameter (PC1 coefficient = 0.51), thickness (0.46), bulk CaCO_3_ content (0.46), peak height (0.43), and higher optimum DIC (0.33). Calcite crystal length of shells (0.13) and width of the Gaussian preference function (-0.01) did not contribute to the separation of the strains.

## Discussion

Climate and land use changes are globally accelerating terrestrial and aquatic carbon cycling rates and thereby modify chemical processes in aquatic systems (de Wit et al. 2023). CO_2_-induced acidification in oceans is predicted to be particularly detrimental for marine shell-forming organism and a diversity of negative and variable responses to calcification between species and strains in marine phytoplankton has been reported (Ries et al. 2009, Müller et al. 2015, Comeau et al. 2017, Page et al. 2022). Responses of freshwater calcifying phytoplankton have received far less attention and are more difficult to predict as effects on bicarbonate and carbonate ion concentrations in lakes operate via complex geochemical pathways (Hasler et al. 2016, Ninokawa and Ries 2022).

Using experimental manipulation of the carbonate system, we have shown that dissolved inorganic carbon concentration appears to be an important driver controlling *P. lenticularis* calcification and shell morphology parameters. The 12 *P. lenticularis* strains responded with a significant reduction in calcification, shell thickness and formation of calcite crystals under declined DIC concentration, while these parameters remained constant at elevated DIC after 14 days of incubation. A decreased calcification and reduction in shell thickness of *P. lenticularis* shells at low DIC concentrations ≤ 1 mmol l^-1^ is likely explained by the algae’s disability to extract sufficient dissolved inorganic carbon from the culture medium and efficiently convert it into calcite, the major constituent of their shells. Declining DIC and pH result in decreased concentration of carbonate ions (CO_3_^2-^) and consequently in a decreased calcite saturation state, on which aquatic calcifiers depend. The final external maturation stage in *P. lenticularis* shell formation may render less control and therefore limits the capacity for adaptation to changing carbonate chemistry. However, we found that *P. lenticularis* has demonstrated a high potential for acclimation and extensive plasticity of individual strains to a wide range of changes in carbonate chemistry showing negative calcification responses only to low DIC levels.

The mineralization pathways in *P. lenticularis* are just being investigated (Shaked et al. 2023). In contrast to coccolith formation, ions are concentrated after endocytosis in *P. lenticularis* and crystals are not fully formed before exiting the cell. These small crystals are then located on the outer surface of the daughter cell lorica, and more crystals form around them until the gaps between the crystals on the outer surface of the lorica are closed. Observations of cells with complete shells show that there likely are additional stages of shell maturation after the reproduction cycle is complete (Hepperle and Krienitz 1996). Fully formed shells can have different shell thicknesses and various levels of organization according to their environment (Lenz et al. 2018). The capability of maintaining elevated pH at the sites of calcification in artificially acidified waters was recently demonstrated across a range of phyla (Gilbert et al. 2022). Studies of single-celled organisms in marine environments demonstrate that control of cytoplasmic pH in coccolithophores involved voltage-gated H^+^ channels in the plasma membrane pushing H^+^ out of the cell in order to maintain saturation conditions for calcite formation (Taylor et al. 2017, Dassow 2022). H^+^ channels showed greatly reduced activity in cells acclimated to low pH which impaired the ability to remove H^+^ generated by the internal calcification process (Kottmeier et al. 2022). The energization of plasma membrane transport in freshwater algae is less clear. *P. lenticularis* belongs to the Chlamydomonadales family of freshwater green algae. An ability to generate electrical voltage differences across their membrane has been found in Chlamydomonas and Characean algae (Merchant et al. 2007, Beilby et al. 2022). There is also evidence suggesting that several freshwater algal taxa utilize a combination of H^+^- and Na^+^-energized transport at the plasma membrane, which may enable them to adapt to changes in the pH and ionic composition of their environments (Taylor et al. 2012). The capacity of pH regulation through voltage-gated proton channels may be central to counter the effects of declining DIC and pH, but needs further investigation in freshwater algae.

In the last decades, studies investigating responses to ocean acidification of *E. huxleyi* observed that high H^+^ concentration in seawater likely correspond with high H^+^ concentration inside phytoplankton cells and lead to reduced CO_2_ fixation efficiency in the chloroplasts and decreased algal growth (Bach et al. 2011). We showed, that *P. lenticularis* growth rates remained stable over a wide range of pH and DIC levels in the culture experiments. *P. lenticularis* seems to be capable of keeping their cell division rate constant over a pH range between approx. 6 to 9 and exponentially increasing carbonate ion concentration up to a DIC of 8 mmol/l. A potential ability to activate and deactivate a CO_2_ concentrating mechanism (CCM) in response to changing CO_2_ and HCO_3_^-^ conditions could explain the acclimation of *P. lenticularis* to variable DICs (Kottmeier et al. 2016). In *Chlamydomonas reinhardtii*, a well studied member of the Chlamydomonadales family, at least 12 genes that encode carbonic anhydrase isoforms, have been detected (Moroney et al. 2011). Carbonic anhydrases (CAs) are zinc-containing metalloenzymes and important components of the CCM. CAs catalyze the reversible interconversion of CO_2_ and HCO_3_^-^. A sudden and entire reduction in growth of most *P. lenticularis strains* at ≥ 10 mmol l^-1^ DIC may be explained by the cells disability to regulate the intracellular pH at high extracellular pH > 9. At the cellular level, a reaction to elevated carbonate concentrations involves regulating of membrane fluidity, metabolism, and photosynthesis. These costly regulatory mechanisms may slow down cell division, while enhancing survival at higher DIC.

The two lakes chosen for isolation of *P. lenticularis* differ substantially in morphology, trophical status and carbonate inputs offering a comparison between strains that were exposed to higher variation in one lake compared to the other. The *P. lenticularis* strains used in this study were isolated a few month prior to the experiment. Adaptation in monoclonal cultures is assumed to be slow due to asexual reproduction and genetic changes are based mainly on mutations. The phenotypic differences observed in the experiments are likely based on genetic or plastic differences the strains established prior to cultivation. Exploring intra-population variability in physiological experiments offers a wider perspective for evaluating species traits and is essential to predict future responses to environmental changes.

Our results demonstrate that *P. lenticularis* strains are adapted to the specific environmental conditions in their lake of origin. Local adaptation occurs when resident individuals have a higher fitness in their local environment than those from other populations of the same species due to genetic change (Pigliucci 2001). In our experiments OS strains developed a clear growth optima at DIC 4 mmol l^-1^, the ambient average DIC level in their lake of origin. Growth optima of GS strains were more variable and tended to shift to higher DIC levels between 5 and 6 mmol l^-1^, the ambient average DIC level lake Gönningersee. Consistent growth optima for specific DIC concentrations by strains from the same lake suggest genetic differentiation of populations into distinct phenotypes and local adaptation.

In comparison to OS strains, GS strains developed higher growth rates and significantly larger as well as heavier calcite shells across a wider DIC gradient. Furthermore, intra-population variability in final cell density was higher in GS strains compared to OS strains. Our results demonstrate the importance of using multiple strains in order to make conclusions regarding species traits. DIC concentrations and pH measured during the time of strain isolation were more dynamic in lake Gönningersee compared to measurements from lake Großer Ostersee (by 2 units). Lake Gönningersee is smaller than lake Großer Ostersee and receives direct input from a tributary that frequently carries high loads of carbonate ions from limestone bedrock in the surrounding watershed. The difference in environmental dynamics in Gönningersee may have caused an increased tolerance to higher variation in DIC in GS strains resulting in their higher tolerance to the experimental DIC manipulations. A cell’s physiological reaction to the current environment is determined by complex interactions between the individuals history of environmental exposure and mechanisms governing its plasticity (Kremer et al. 2018). Local adaptation in phytoplankton occurs by lineage selection on new arrivals, or by natural selection on standing genetic variation leading to selection of individuals best adapted to the local environment (Orsini et al. 2013). *P. lenticularis* forms resting stages, which may act as a genetic backup in the sediment and buffer against new arrivals. Schaum et al. 2013 have shown that phytoplankton populations from more variable environments are more tolerant to a changing environment. We can confirm these findings with our experimental results on *P. lenticularis* strains. The exact mechanisms underlying adaptation and genetic composition in freshwater calcifiers remain an important topic of study.

Freshwater habitats are often isolated water bodies with large variation in environmental variables that provide stronger dispersal barriers and potential for differentiation than marine habitats. The functional diversity of freshwater calcifying algae may be considerably larger than is presently assumed based upon only a limited number of studies and strains. The ability of *P. lenticularis* to continue its contribution to autochthonous calcite precipitated in the surface layer of lakes and ponds will depend on their ability to survive shifts in the composition and function forecasted for freshwater lakes over the next centuries (Capon et al. 2023). Individual variability will maintain high functional diversity in *P. lenticularis* populations and increases success to persist in a changing environment.

## Conclusion

The present study shows that calcification and shell morphology of *P. lenticularis* is sensitive to decreasing DIC levels from 4 to 1 mmol l^-1^ during 14 d of incubation. Artificial reduction of DIC levels resulted in a significant decrease in shell thickness, loss of a distinct crystalline structure at the cell surface and decrease in cell diameter. Ambient crystal length and shell thickness was preserved at increasing DIC from 4 to 8 mmol l^-1^, but crystals either elongated to spines or developed an extensive rhombohedral shape. We did not find changes in growth rates as a function of the DIC manipulation. However, cultures that grew at low and high DIC levels showed reduced cell densities at day 14. Cell densities followed a Gaussian response profile with highest densities at ambient DIC concentrations measured in their lake of origin. This laboratory experiment using freshly isolated genotypes from two lakes examines biogeochemical processes through the cellular perspective and paves the road for further studies of physiological, metabolic and molecular responses of calcifying freshwater algae to predict future freshwater carbon cycles.

## Acknowledgements

We thank Maren Lentz and Sebastian Lenz for laboratory assistance during the experiments. The help and advice from Lothar Krienitz concerning the strain isolation and experimental set-up of the DIC experiment was very appreciated. We gratefully acknowledge the Tübingen Structural Microscopy Core Facility (funded by the Excellence Strategy of the German Federal and State Governments) for their support and assistance in this work. We thank two anonymous reviewers whose comments greatly improved this paper.

